# Seeing spots: Quantifying mother-offspring similarity and assessing fitness consequences of coat pattern traits in a wild population of giraffes (*Giraffa camelopardalis*)

**DOI:** 10.1101/161281

**Authors:** Derek E. Lee, Douglas R. Cavener, Monica L. Bond

## Abstract

Polymorphic phenotypes of mammalian coat coloration have been important to the study of genetics and evolution, but less is known about the inheritance and fitness consequences of individual variation in complex coat pattern traits such as spots and stripes. Giraffe coat markings are highly complex and variable and it has been hypothesized that variation in coat patterns most likely affects fitness by camouflaging neonates against visually hunting predators. We quantified complex coat pattern traits of wild Masai giraffes using image analysis software, determined the similarity of spot pattern traits between mother and offspring, and assessed whether variation in spot pattern traits was related to fitness as measured by juvenile survival. The methods we described could comprise a framework for objective quantification of complex mammal coat pattern traits based on photographic coat pattern data. We demonstrated that some characteristics of giraffe coat spot shape were likely to be heritable, as measured by mother-offspring regression. We found significant variation in juvenile survival among phenotypic groups of neonates defined by multivariate clustering based on spot trait measurement variables. We also found significant variation in neonatal survival associated with spot size and shape covariates. Spot trait variation also may be relevant to other components of fitness, such as adult survival or fecundity. These findings will inform investigations into developmental and genetic architecture of complex mammal coat patterns and their adaptive value.

## Introduction

Complex color patterns such as spots and stripes are found on many animal species and these phenotypic traits are hypothesized to play adaptive roles in predator and parasite evasion, thermoregulation, and communication (Cott, 1940; Caro, 2005). Many foundational studies of coloration using starkly different color morphs from diverse taxa such as insects (Kettlewell, 1955; Wittkopp et al., 2003), mice (Morse, 1978; Russell, 1985; Bennett & Lamoreux, 2003), reptiles (Rosenblum et al., 2004; Calsbeek et al., 2008), fish (Endler, 1983; Irion et al., 2016), and birds (Roulin, 2004) demonstrated Mendelian inheritance and natural selection, and discovered genes that cause color morph mutations (Hoekstra, 2006; Protas & Patel, 2008; San-Jose & Roulin, 2017). Individual variation in a complex color pattern trait of spot size was also part of the earliest work on genetics and inheritance (Wright, 1917). Measuring individual variation in complex color patterns, especially detailed measurements such as animal biometrics (Kuhl & Burghardt, 2013), can provide novel insight into developmental and genetic architecture (Bowen & Dawson, 1977; Klingenburg, 2010; San-Jose & Roulin, 2017), and the adaptive value of the patterns (Hoekstra, 2006; Allen et al., 2010), as well as benefitting studies of behavior (Lorenz, 1937; Whitehead, 1990), and population biology (Holmberg et al., 2009; Lee & Bolger, 2017), and the growing field of phenomics (Houle et al., 2010). Some methods to robustly quantify individuals’ continuous variation in complex color patterns have been developed for general use (Schneider et al., 2012; Van Belleghem et al., 2018) and specific taxa such as fishes (Endler, 1980; Holmberg et al., 2009), butterflies (LePoul et al., 2014), penguins (Sherley et al., 2010), and primates (Allen et al., 2015). We see a need for more tools and techniques to reliably quantify individual variation in complex coat pattern traits in wild populations (Eizirik et al., 2010; Willisch et al., 2013), and studies that use quantitative genetics and demographic methods to investigate heritability and adaptive significance of those traits in wild mammal populations (Kruuk et al., 2008; Kaelin et al., 2012).

The coat patterns of Masai giraffes (*Giraffa camelopardalis tippelskirchii*) are complex and show a high degree of individual variation (Dagg, 1968; **Figure 1**). Masai giraffes spots vary in color and shape from those that are nearly round with very smooth edges (low tortuousness), to extremely elliptical with incised or lobate edges (high tortuousness). Giraffe skin pigmentation is uniformly dark grey (Dimond & Montagna, 1976), but the spots that make up their coat markings are highly variable in traits such as color, roundness, and perimeter tortuousness. This variation has been used to classify subspecies (Lydekker, 1904), and to reliably identify individuals because patterns do not change with age (Foster, 1966; Bolger et al., 2012; Dagg, 2014). Dagg (1968) first presented evidence from a small zoo population that the shape, number, area, and color of spots in giraffe coat patterns may be heritable, but analysis of spot traits in wild giraffes, and objective measurements of spot characteristics in general have been lacking.

**Figure 1.**
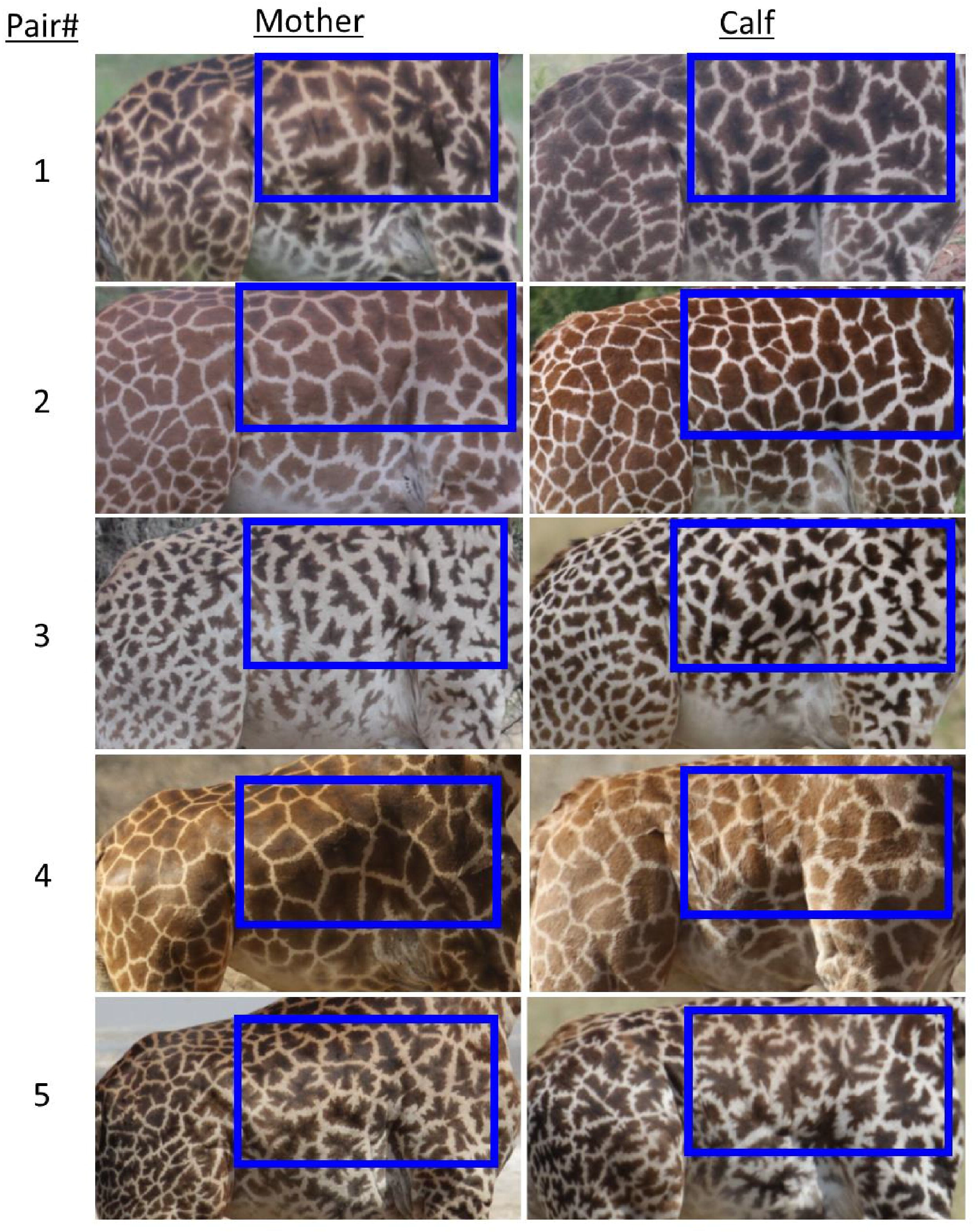
Representative images of spot patterns of mother-calf pairs of Masai giraffes **(*Giraffa camelopardalis tippelskirchii*) from the Tarangire ecosystem, Tanzania used in this study.**

The blue rectangle shows the area analysed using ImageJ to characterize spot pattern traits.

It has been hypothesized that giraffe coat patterns evolved to camouflage neonates whose primary defense against predation is concealment (Langman, 1977; Mitchell & Skinner, 2003); thus the most likely fitness effects from variation in coat patterns should be variation in juvenile survival. Giraffe calves spend much of their time, day and night, hiding in the dappled light of trees and bushes and their ability to match this background should influence detection by visually hunting predators such as lions and hyenas (Endler, 1978; Merilaita et al., 2017). Background matching, the adaptation of an animal’s coloration to mimic its average background and reduce detection by visually hunting predators, is a common form of camouflage (Endler, 1978; Merilaita et al., 2017). Alternative hypotheses about the adaptive value of giraffe coat markings include thermoregulation (Skinner & Smithers, 1990), and in this social species with good visual sensory perception (Dagg, 2014; VanderWaal et al., 2014), markings could also facilitate individual recognition (Tibbetts & Dale, 2007) and kin recognition (Beecher, 1982; Tang-Martinez, 2001).

Our purpose in this study was to: 1) demonstrate the use of public domain image analysis software ImageJ (Schneider et al., 2012) to extract patterns from image data and quantify multiple aspects of the complex coat patterns of wild Masai giraffes; 2) use quantitative genetics methods (parent-offspring regression) to quantify the proportion of observed phenotypic variation of a trait that is shared between mother and offspring; and 3) determine whether variation in complex coat pattern traits was related to a measure of fitness (survival) and thereby infer the effect of natural selection (viability selection) on giraffe coat patterns (Lande & Arnold, 1983; Falconer & Mackay, 1996).

## Materials & Methods

As a general overview, our methods were to: 1) collect field data in one area of Tanzania as digital images of giraffes to be used for spot pattern and survival analyses; 2) extract patterns from images; 3) quantify giraffe patterns by measuring 11 spot traits; 4) use principal components analysis (PCA) to reduce the dimensionality of the spot traits; 5) use mother-offspring regressions to estimate the phenotypic similarity between mother and offspring of the 11 spot traits and the 1st two dimensions of the PCA; 6) use k-means clustering to assign giraffe calves into 4 phenotypic groups according to their spot pattern traits; 7) use capture-mark-recapture analysis to estimate survival and determine whether there are fitness differences among the phenotypic groups; 8) use capture-mark-recapture analysis to determine whether there are fitness effects from any particular spot traits.

### Field Data Collection

This study used data from individually identified, wild, free-ranging Masai giraffes in a 1700 km^2^ sampled area within a 4400 km^2^ region of the Tarangire Ecosystem, northern Tanzania, East Africa. Data were collected as previously described in Lee et al. (2016a). We collected data during systematic road transect sampling for photographic capture-mark-recapture (PCMR). We conducted 26 daytime surveys for giraffe PCMR data between January 2012 and February 2016. We sampled giraffes three times per year around 1 February, 1 June, and 1 October near the end of every precipitation season (short rains, long rains, and dry, respectively) by driving a network of fixed-route transects on single-lane dirt tracks in the study area. We surveyed according to Pollock’s robust design sampling framework (Pollock, 1982; Kendall et al., 1995), with three occasions per year. Each sampling occasion was composed of two sampling events during which we surveyed all transects in the study area with only a few days interval between events. Each sampling occasion was separated by a 4-month interval (4.3 years × 3 occasions year^-1^ × 2 events occasion^-1^ = 26 survey events).

During PCMR sampling events, a sample of individuals were encountered and either ‘sighted’ or ‘resighted’ by slowly approaching and photographing the animal’s right side at a perpendicular angle (Canon 40D and Rebel T2i cameras with Canon Ultrasonic IS 100 – 400 mm lens, Canon U.S.A., Inc., One Canon Park, Melville, New York, 11747, USA). We identified individual giraffes using their unique and unchanging coat patterns (Foster, 1966; Dagg, 2014) with the aid of pattern-recognition software Wild-ID (Bolger et al., 2012). We attempted to photograph every giraffe encountered, and recorded sex and age class based on physical characteristics. We assigned giraffes to one of four age classes for each observation based on the species’ life history characteristics and our sampling design: neonate calf (0 – 3 months old), older calf (4 – 11 months old), subadult (1 – 3 years old for females, 1 – 6 years old for males), or adult (> 3 years for females, > 6 years for males) using a suite of physical characteristics (Strauss et al., 2015), and size measured with photogrammetry (Lee et al., 2016a). In this analysis, we used only adult females and animals first sighted as neonate calves.

All animal work was conducted according to relevant national and international guidelines. This research was carried out with permission from the Tanzania Commission for Science and Technology (COSTECH) Research Permit numbers 2017-163-ER-90-172, 2016-146-ER-2001-31, 2015-22-ER-90-172, 2014-53-ER-90-172, 2013-103-ER-90-172, 2012-175-ER-90-172, 2011-106-NA-90-172, Tanzania National Parks (TANAPA), the Tanzania Wildlife Research Institute (TAWIRI). No Institutional Animal Care and Use Committee (IACUC) approval was necessary because animal subjects were observed without disturbance or physical contact of any kind.

### Quantification of Spot Patterns

We extracted patterns and analysed spot traits of each animal within the shoulder and rib area by cropping all images to an analysis rectangle that fit horizontally between the anterior edge of the rear leg and the chest, and vertically between the back and where the skin folded beneath the posterior edge of the foreleg (**Figure 1**). For color trait analysis, we used the Color Histogram procedure of ImageJ (Schneider et al., 2012) on full-color images of the analysis rectangle. We extracted coat patterns using ImageJ to convert full-color images of the analysis rectangle to 8-bit greyscale images, then converted to bicolor (black and white) using the Enhance Contrast and Threshold commands (Schneider et al., 2012). We quantified 10 spot trait measurements of each animal’s extracted coat pattern using the Analyze Particles command in ImageJ (Schneider et al., 2012). To account for differences in image resolution and animal size (including age-related growth), and to obtain approximately scale-invariant standard images of each animal, we set the measurement unit of each image equal to the number of pixels in the height of the analysis rectangle. Therefore all measurements are in giraffe units (GU), where 1 GU = height of the analysis rectangle (**Figure 1**). We excluded spots cut off by the edge of the analysis rectangle to avoid the influence of incomplete spots, and we also excluded spots whose area was < 0.00001 GU^2^ to eliminate the influence of speckles.

We characterized each animal’s coat spot pattern traits within the analysis rectangle using the following 11 metrics available in ImageJ: number of spots; mean spot size (area); mean spot perimeter; mean angle between the primary axis of an ellipse fit over the spot and the x-axis of the image; mean circularity (*4*π × *[Area]* / *[Perimeter]^2^* with a value of 1.0 indicating a perfect circle and smaller values indicating an increasingly elongated shape); mean maximum caliper (the longest distance between any two points along the spot boundary, also known as Feret diameter); mean Feret angle (the angle [0 to 180 degrees] of the maximum caliper); mean aspect ratio (of the spot’s fitted ellipse); mean roundness (4 × [*Area*]π × [*Major axis*]^2^ or the inverse of aspect ratio); mean solidity ([*Area*] / [*Convex area*], also called tortuousness); and mode shade ([65536 × r] + [256 × g] + [b] using RGB values from color histogram from full color photos).

We quantified among-individual variation in spot trait values by reporting the mean, SD, and coefficient of variation (CV) of each trait. We also quantified the reliability of our spot pattern trait measurement technique by computing the amount of among-measurement variation for the same animal made on different photos from different dates using a set of 30 animals with >2 images per animal. We performed a principal components analysis (PCA; Hotelling, 1933) on the covariance matrix of the 10 spot trait measurements (standardized) to examine the patterns of variation and covariation among the spot measurement data and to compute 2 summary dimensions explaining the 10 measurements. We performed k-means clustering to divide animals into ‘coat pattern phenotypes,’ phenotypic groups based upon their spot trait characteristics (MacQueen, 1967). The optimal number of phenotypic groups was determined by the gap statistic (Tibshirani et al., 2001). We performed all statistical operations using R (R Core Development Team, 2017).

### Mother-Offspring Similarity of Spot Traits

The (narrow sense) heritability of a trait (symbolized *h^2^*) is the proportion of its total phenotypic variance that is additive, or available for selection to act upon. Parent-offspring (PO) regression is one of the traditional quantitative genetics tools used to test for heritable additive genetic variation (Falconer & Mackay, 1996). PO regression studies cannot distinguish among phenotypic similarity due to genetic heritability, maternal investment, or shared environmental effects, it is however one of the few methods available when information on other kin relations are lacking. Pigmentation traits in mammals are known to have a strong genetic basis (Bennett and Lamoreux 2003; Hoekstra 2006), supporting the interpretation of PO regression as indicating a genetic component. We expect minimal non-random variation due to environmental effects because the calves were all born in the same area with the same vegetation communities during a relatively short time period of average climate and weather with no spatial segregation by coat pattern phenotype (**Supplementary Material Figure S1**). The animal model was not an improvement because we do not know fathers, and we had no known siblings in our dataset, therefore PO regression is the most appropriate tool for our estimates of heritability, with the caveat that there are potentially environmental and maternal investment effects also present.

We identified 31 mother-calf pairs by observing extended suckling behavior. Wild female giraffes very rarely suckle a calf that is not their own (Pratt and Anderson 1979). We examined all identification photographs for individuals in known mother-calf pairs, and selected the best-quality photograph for each animal based on focus, clarity, perpendicularity to the camera, and unobstructed view of the torso.

We predicted spot pattern traits of a calf would be correlated with those of its mother. We estimated the mother-offspring similarity for each of the 11 spot trait measurements, and the first dimension generated by the PCA. When we examined the 11 individual spot traits, we used the Bonferroni adjustment (α/number of tests) to account for multiple tests and set our adjusted α = 0.0045. We performed statistical operations in R (R Core Development Team, 2017).

### Fitness of Spot Patterns from Juvenile Survival

We assembled encounter histories for 258 calves first observed as neonates for survival analysis. For each calf we selected the best-quality calf-age (age < 6 mo) photograph based on focus, clarity, perpendicularity to the camera, and unobstructed view of the torso, and ran the photographs through the ImageJ analysis to quantify each individual’s coat spot traits. We analysed survival using capture-mark-recapture apparent survival models that account for imperfect detectability during surveys (White & Burnham, 1999). No capture-mark-recapture analyses except ‘known fate’ models can discriminate between mortality and permanent emigration, therefore when we speak of survival it is technically ‘apparent survival,’ but during the first seasons of life we expected very few calves to emigrate from the study area, and if any did emigrate permanently this effect on apparent survival should be random relative to their spot pattern characteristics.

We estimated age-specific seasonal (4-month seasons) survival (up to 3 years old) according to coat pattern phenotype groups with calves assigned to groups by k-means clustering of their overall spot traits. We compared 3 models, a null model of no group effect, age + group, and age × group, to examine whether coat pattern phenotypes affected survival differently at different ages. We also estimated survival as a function of individual covariates of specific spot traits including linear and quadratic relationships of all 11 spot traits and the first two PCA dimensions on juvenile survival to examine whether directional, disruptive, or stabilizing selection was occurring (Lande & Arnold, 1983; Falconer & Mackay, 1996). To determine at what age specific spot traits had the greatest effect of survival, we examined survival as a function of spot traits during 3 age periods: the first season of life, first year of life, and first three years of life.

We used Program MARK to analyse complete capture-mark-recapture encounter histories of giraffes first sighted as neonates (White & Burnham, 1999). We analysed our encounter histories using Pollock’s Robust Design models to estimate age-specific survival (Pollock, 1982; Kendall et al., 1995), and ranked models using AICc following Burnham and Anderson (2002). We used weights (W) and likelihood ratio tests as the metrics for the strength of evidence supporting a given model as the best description of the data (Burnham & Anderson, 2002). Due to model selection uncertainty, we present model-averaged parameter values and based all inferences on these model-averaged values (Burnham & Anderson, 2002). We considered factors to be statistically significant if the 85% confidence interval of the beta coefficient did not include zero (Arnold, 2010), this is a widely accepted convention for covariates of survival analyses (Arnold, 2010).

Based on previous analyses for this population (Lee et al., 2016a, b), we constrained parameters for survival (S) and temporary emigration (γ□ and γ□) to be linear functions of age (symbolized ‘A’), and capture and recapture (c and p) were time dependent (symbolized ‘t’), so the full model was: {(S(A), γ□(A), γ□(A), c(t), p(t)}. Giraffe calf survival does not vary by sex (Lee et al., 2016b), so we analysed all calves together as an additional constraint on the number of parameters estimated. We tested goodness-of-fit in encounter history data using U-CARE (Choquet et al., 2009), and we found some evidence for lack of fit (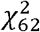 = 97, *P* = 0.01), but because the computed ̂ adjustment was < 3 (̂ = 1.5), we felt our models fit the data adequately and we did not apply a variance inflation factor (Burnham and Anderson 2002; Choquet et al., 2009).

We have deposited the primary data underlying these analyses as follows: Sampling locations, original data photos, and spot trait data: Dryad DOI: https://doi.org/10.5061/dryad.6514r.

## Results

We were able to extract patterns and quantify 11 spot traits using ImageJ, and found measurements were highly reliable with low variation in measurements from different photos of the same individual (**Table 1**). From our 31 mother-calf pairs, we found two spot traits, circularity and solidity (tortuousness) (**Figure 2**) had significant PO slope coefficients between calves and their mothers indicating similarity (**Table 1**). Neither of the first two dimensions from the PCA (see below) had significant PO regression slopes.

**Table 1.**
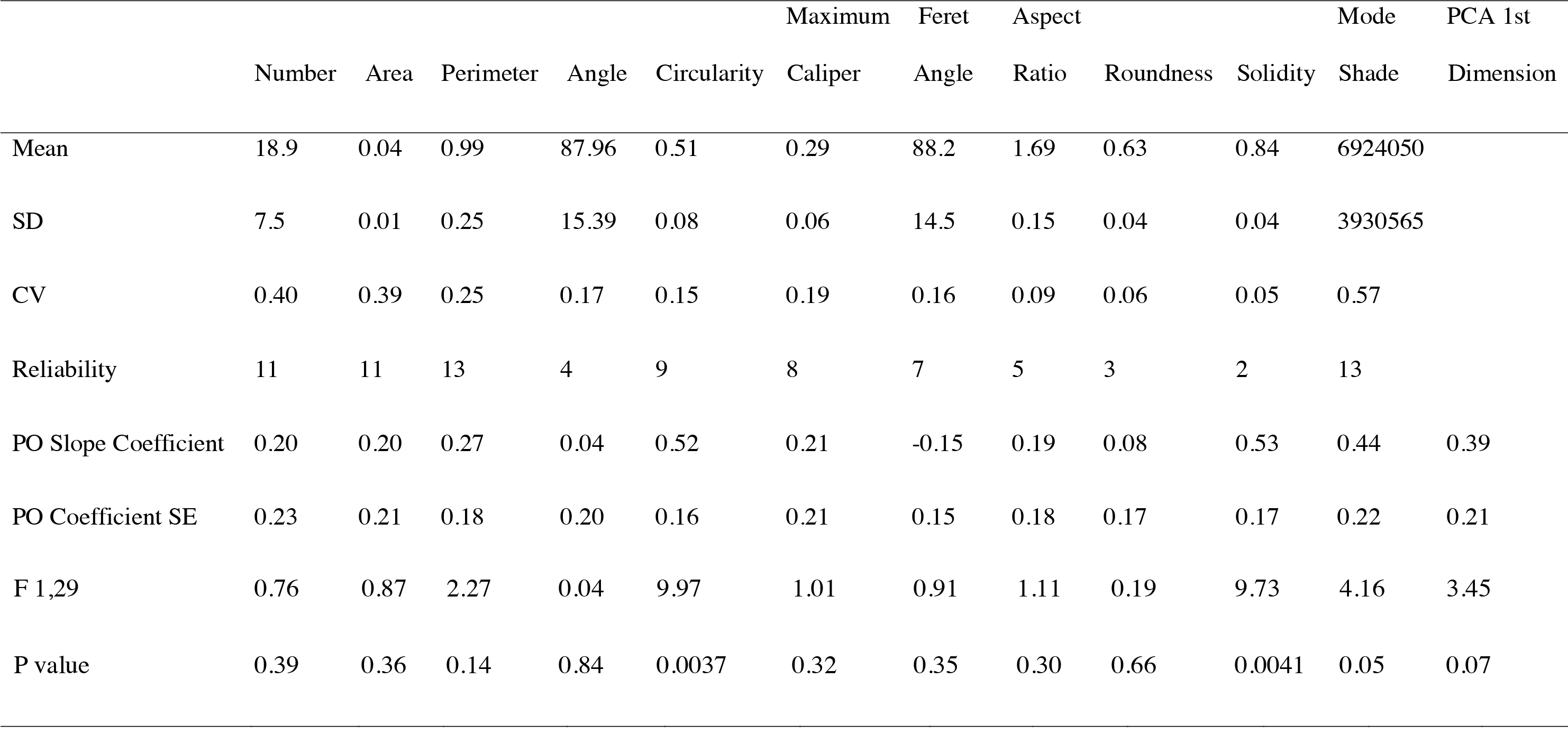
Summary statistics for mother-offspring regressions of spot traits of Masai giraffes in northern Tanzania. Mean trait values, SD (standard deviation), CV (among-individuals coefficient of variation), Reliability (mean % variation in measurements from different pictures of the same individual), PO slope coefficients, F-statistics, and P values are provided. Statistically significant heritable traits are in bold.

**Figure 2.**
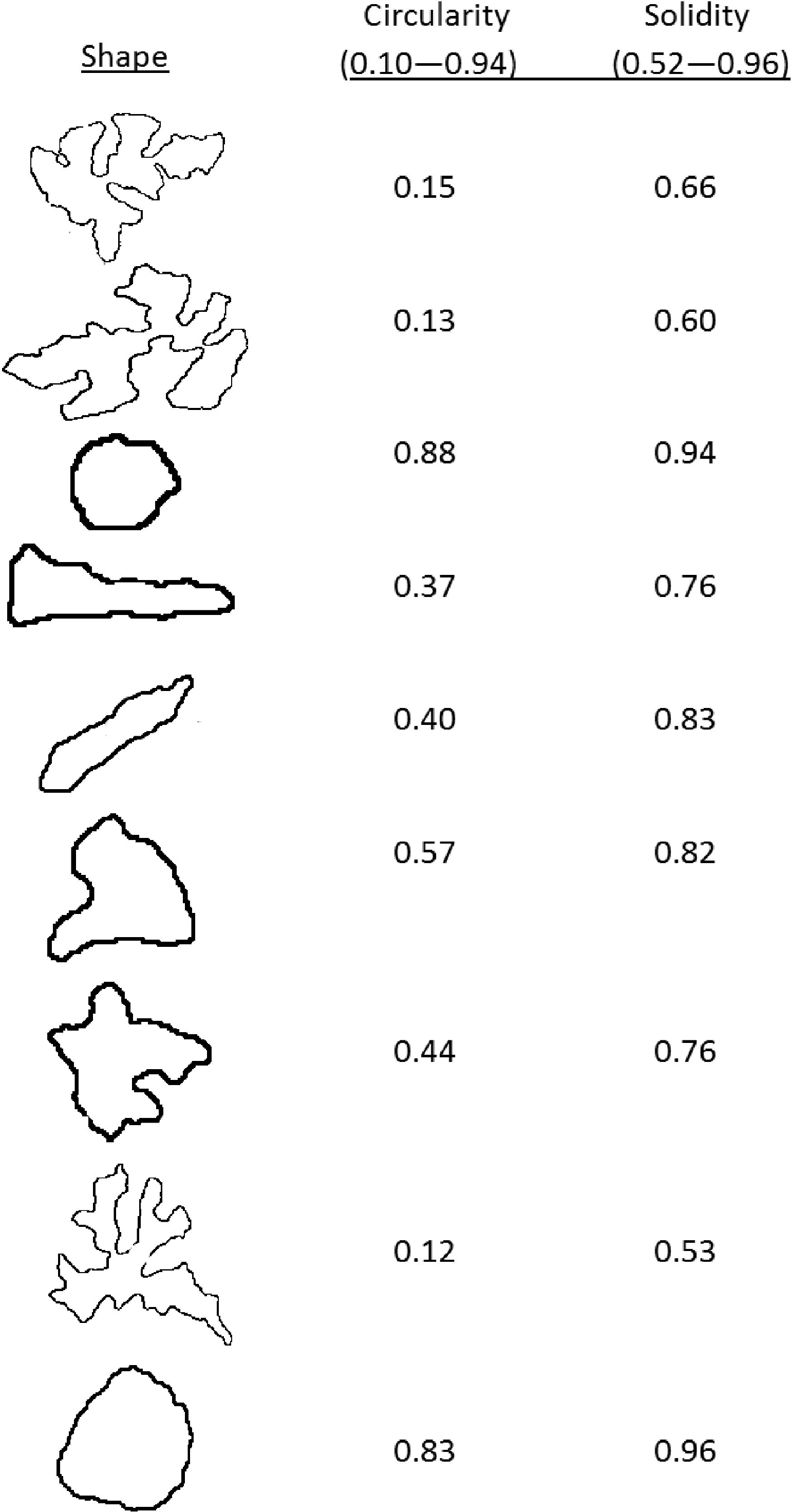
Representative spot outlines from Masai giraffes in northern Tanzania and their corresponding circularity and solidity values.

Ranges of spot trait values from 258 calves are given in parentheses.

The first dimension from the PCA (from 258 calves, including the 31 calves used to estimate heritability) was composed primarily of spot size-related traits (perimeter, maximum caliper, area, and number) such that increasing dimension 1 meant increasing spot size. Dimension 1 explained 40.5% of the variance in the data (**Figure 3**). The second dimension was composed primarily of spot shape traits (aspect ratio, roundness, solidity, and circularity) such that increasing dimension 2 meant increasing roundness and circularity while decreasing dimension 2 meant more tortuous edges and irregular shapes. Dimension 2 explained 24.0% of the variation in the data (**Figure 3**).

**Figure 3.**
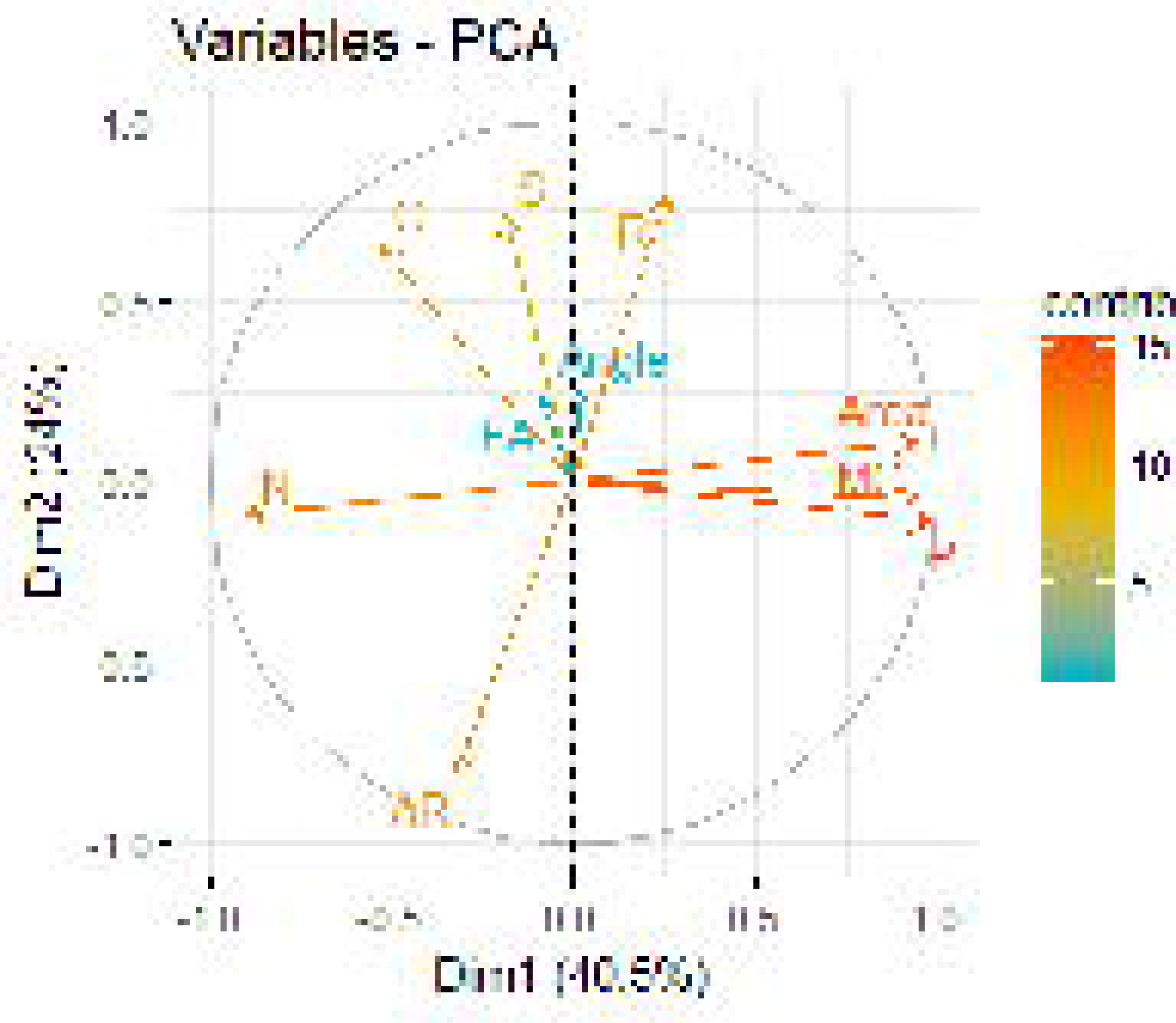
Contributions of 10 trait measurement variables to the first 2 dimensions of the principal components analysis of giraffe spots.

The first dimension (Dim1) was composed primarily of spot size-related traits (perimeter, maximum caliper, area, and number of spots), the second dimension (Dim2) was composed primarily of spot shape traits (aspect ratio, roundness, solidity, and circularity).

Gap statistics indicated 4 phenotypic groups was the optimal number of clusters for k-means clustering, but groups 1 and 2 had a large amount of overlap in PCA variable space (**Figure 4**), so we also defined 3 phenotypic groups by lumping the two overlapping groups. Group 1 had medium-sized circular spots, group 2 had small-sized circular and irregular spots, group 3 had medium-sized irregular spots, and group 4 had large circular and irregular spots (**Figures 3** & **4**). Our survival analysis of 258 calves divided into 4 phenotypic groups based on their spot traits indicated that the null model was top-ranked, but AIC_c_ weights showed there was some evidence for survival variation among the 4 phenotypic groups (**Table 2**). The 3 phenotypic group model found significant differences in survival according to group (**Table 2**, beta coefficient for lumped groups 1 and 2 = −0.717, 85% CI = −1.235 to −0.199). Model-averaged seasonal apparent survival estimates indicated differences in survival existed among phenotypic groups during the first seasons of life, but those differences were greatly reduced in ages 1 and 2 years old (**Figure 5**).

**Figure 4.**
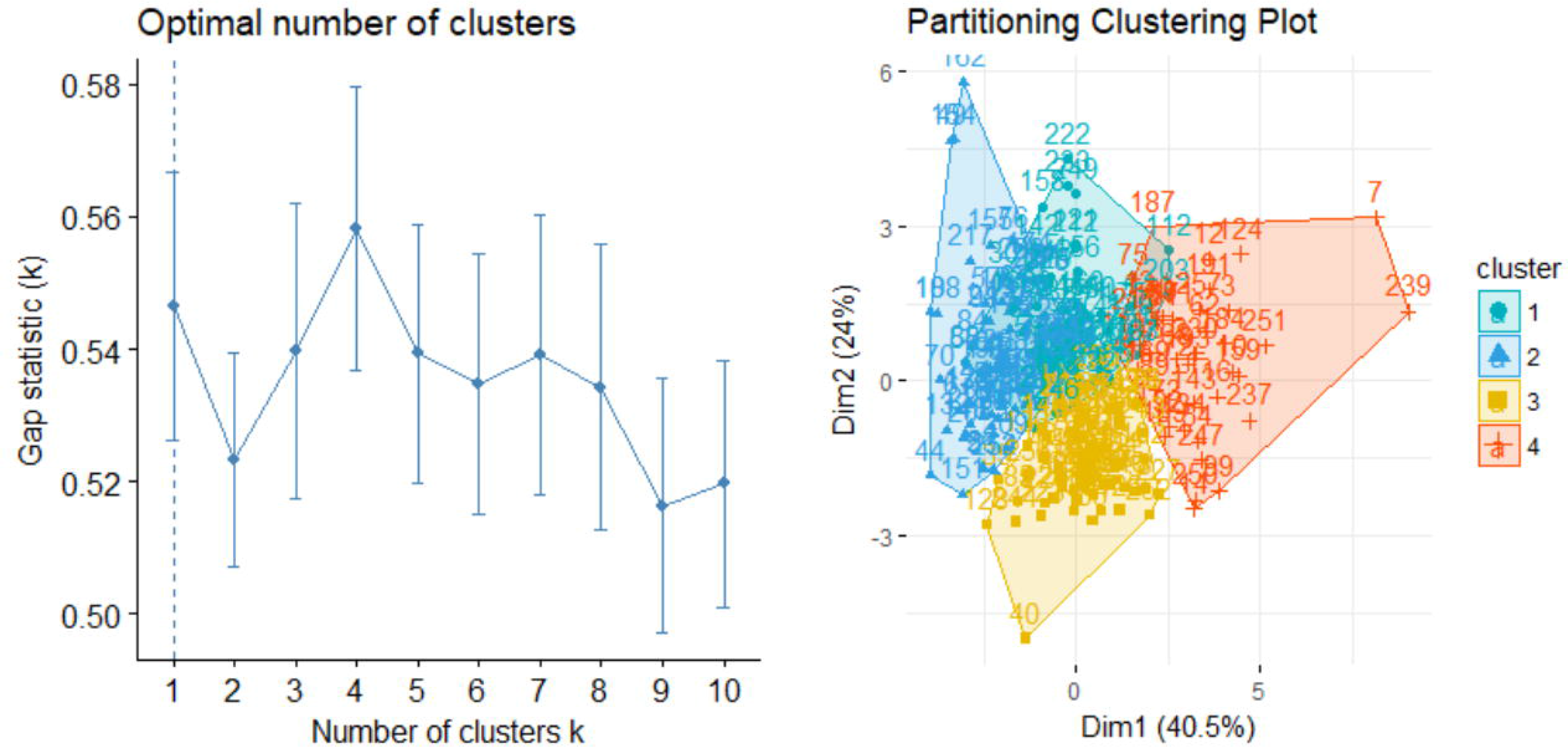
Results from k-means cluster analysis of giraffe spot patterns to define phenotypic groups.

Left is gap statistic for different numbers of groups. Right is 4 clusters mapped in PCA space.

**Figure 5.**
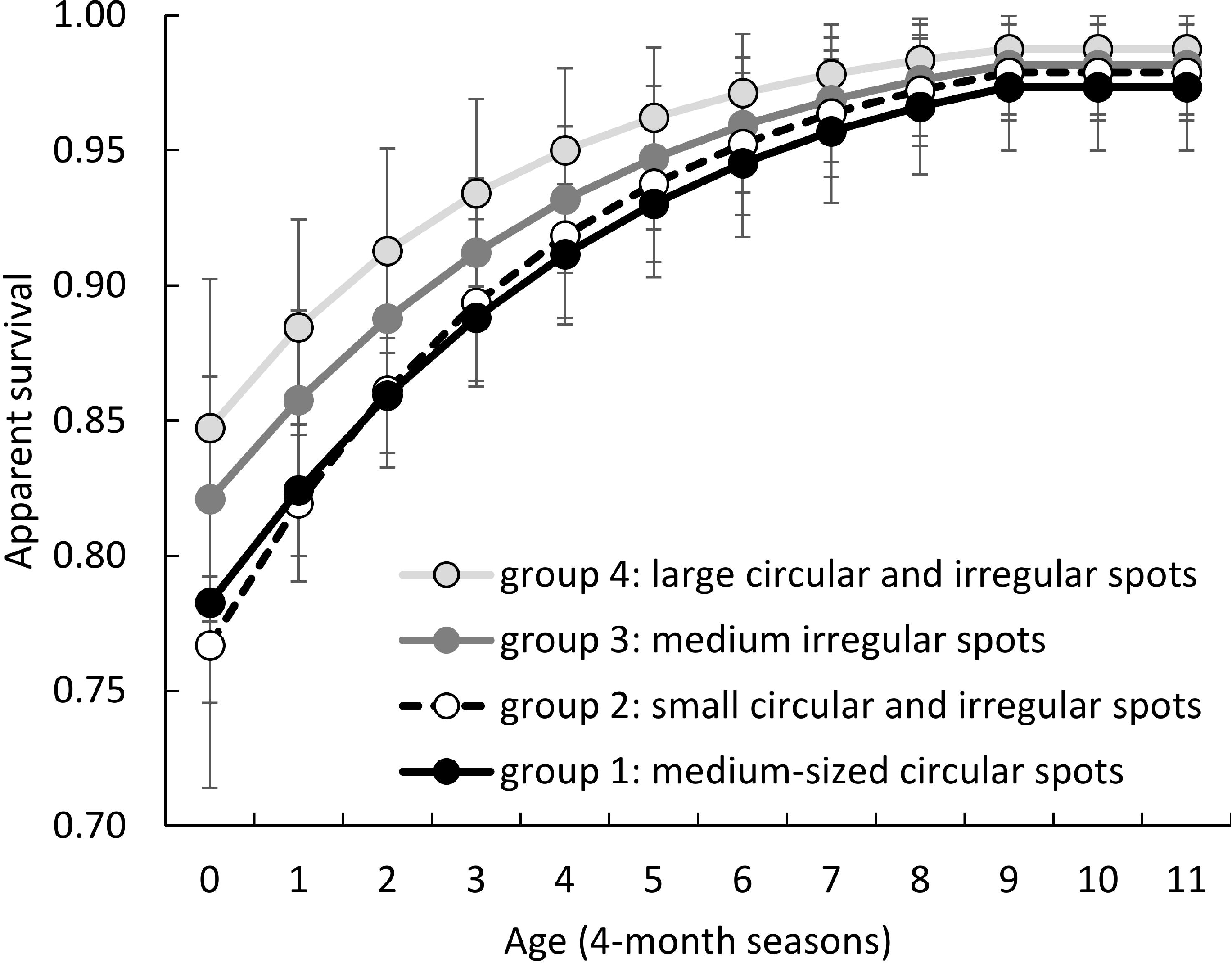
Model-averaged seasonal (4 months) apparent survival estimates for coat pattern phenotypic groups of giraffes defined by k-means clustering of their spot pattern traits.

There was evidence for significant differences in survival among phenotypic groups during the younger ages, but those differences were greatly reduced as the animals approached adulthood (age 9 – 11 seasons). Error bars are ±1 SE.

**Table 2.** Model selection results for giraffe calf survival according to phenotypic groups defined by spot traits. Model weights indicated some evidence for group effects on survival. Notation ’A’ indicates a linear trend with age. Additive models indicate groups shared a common slope coefficient, but had different intercepts; multiplicative models indicated groups had different intercepts and different slopes. Model structure in all cases was {S(A …) g”(A) g’(A) p(t) c(t)}. Minimum AICc = 3236.38, W = AIC_c_ weight, k = number of parameters.

| Model | $\Delta AICc$ | W | k |
| --- | --- | --- | --- |
| A + 3 groups | 0 | 0.43 | 36 |
| Null (no group effect) | 0.94 | 0.27 | 34 |
| A + 4 groups | 2.06 | 0.15 | 37 |
| A × 4 groups | 3.01 | 0.09 | 40 |
| A × 3 groups | 3.91 | 0.06 | 38 |

We found two specific spot traits significantly affected survival during the first season of life (number of spots and aspect ratio; beta number of spots = −0.031, 85% CI = −0.052 to −0.009; beta aspect ratio = −0.466, 85% CI = −0.827 to −0.105). Both number of spots and aspect ratio were negatively correlated with survival during the first season of life (**Figure 6**). No other trait during any age period significantly affected juvenile survival (all beta coefficient 85% CIs included zero), but model selection uncertainty was high (**Table 3**).

**Figure 6.**
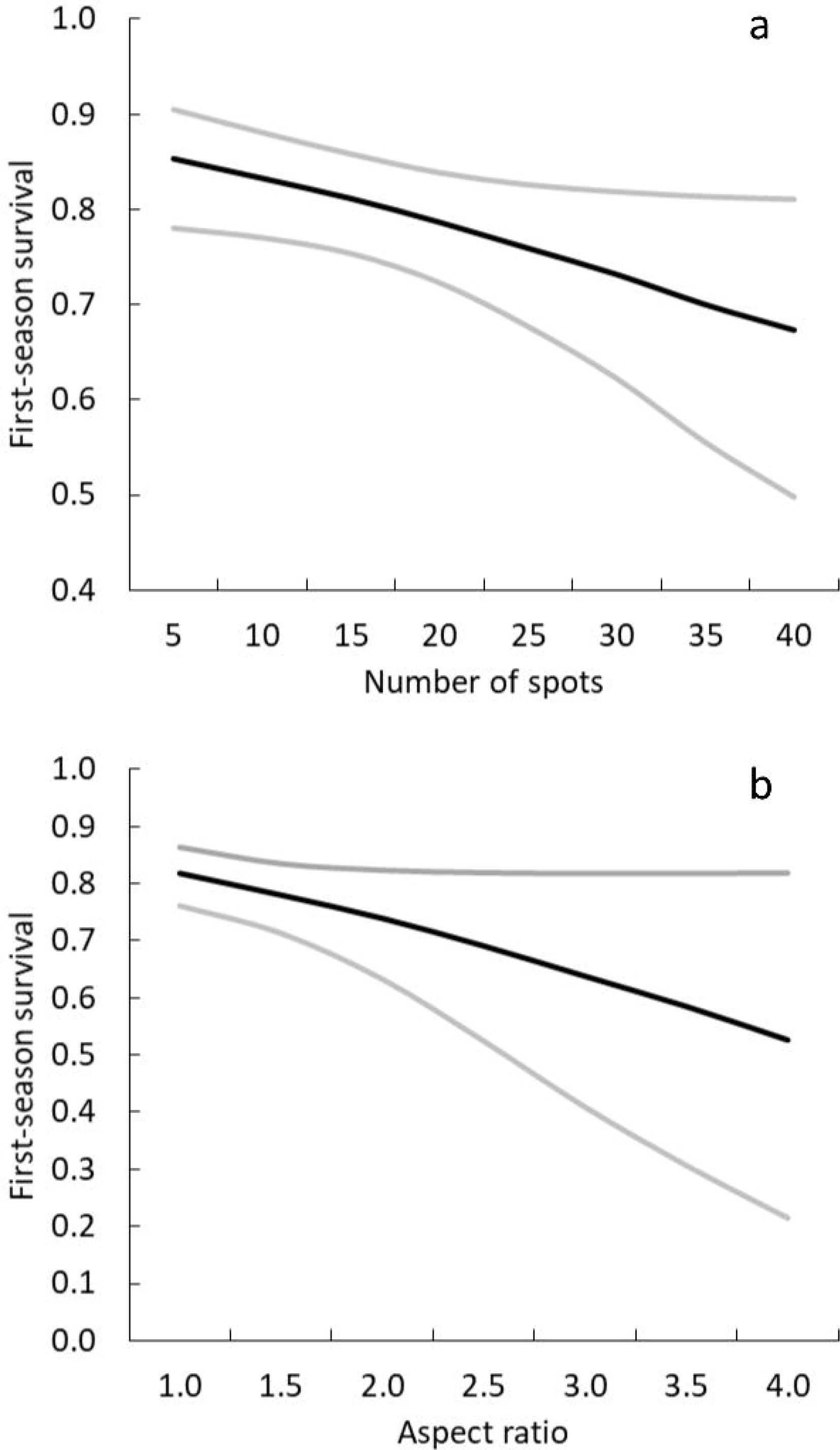
Survival of neonatal giraffes during their first 4-months of life was negatively correlated with number of spots (a) and aspect ratio (b).

Number of spots and aspect ratio are the inversely related to spot size and roundness (the variables used when describing coat pattern phenotypic groups), respectively. Black lines are model estimates, grey lines are 95% confidence intervals.

**Table 3.**
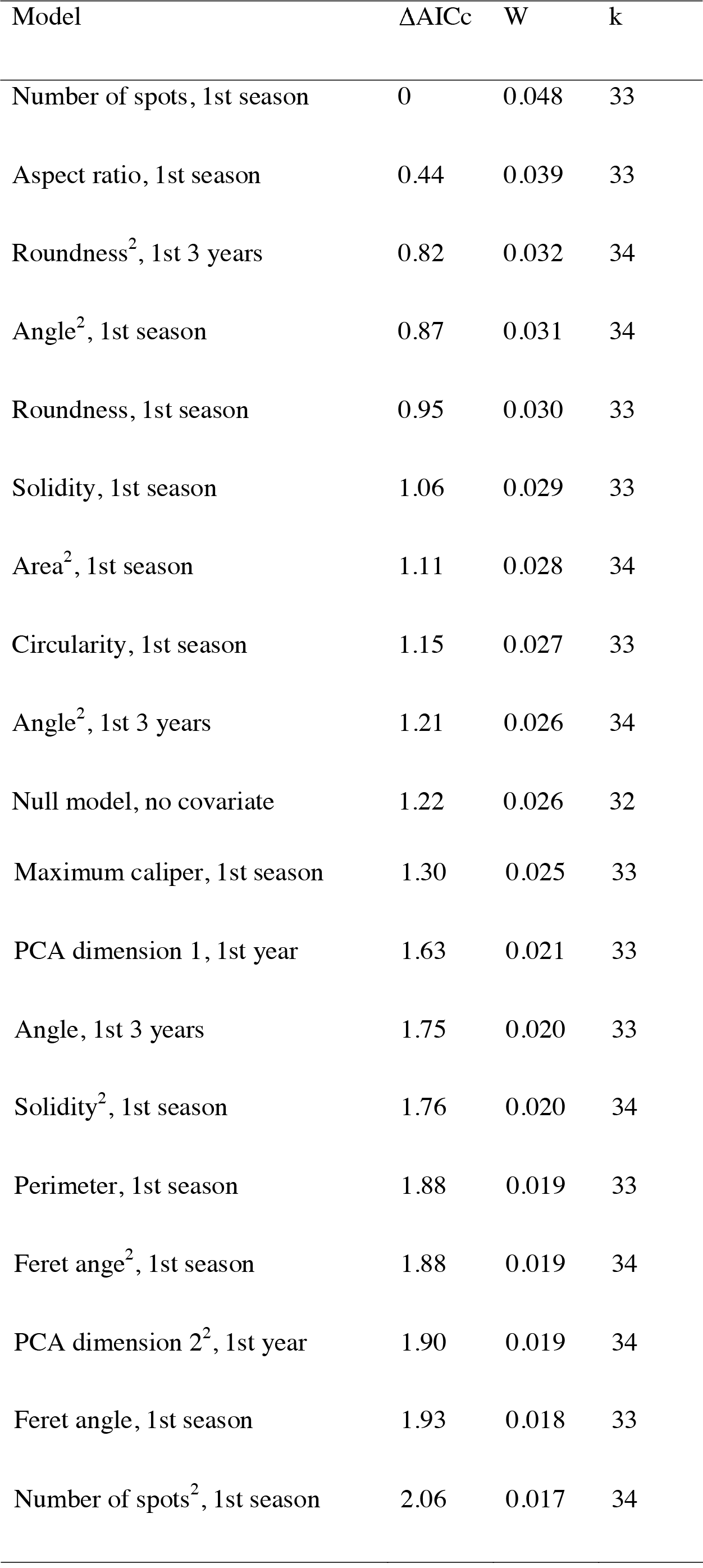
Model selection results for giraffe calf survival as a linear or quadratic function of spot trait covariates during the first season (4 months), first year, and first 3 years of life. Confidence intervals of beta coefficients for two traits excluded zero (number of spots, and aspect ratio), indicating evidence for significant spot trait effects on calf survival during the first season of life. Model structure in all cases was {S(A + Covariate) g”(A) g’(A) p(t) c(t)} with covariate structure in survival. Notation ’A’ indicates a linear trend with age, ’t’ indicates time dependence. Minimum AICc = 3239.87, W = AICc weight, k = number of parameters. Models comprising the top 50% cumulative W are shown.

## Discussion

We were able to objectively and reliably quantify coat pattern traits of wild giraffes using image analysis software. We demonstrated that some giraffe coat pattern traits of spot shape appeared to be heritable from mother to calf, and that coat pattern phenotypes defined by spot size and shape differed in fitness as measured by neonatal survival. Individual covariates of spot size and shape significantly affected survival during the first 4 months of life. These results support the hypothesis that giraffe spot patterns are heritable (Dagg, 1968), and affect neonatal calf survival (Langman, 1977; Mitchell & Skinner, 2003). The fact that spot patterns affected survival could be related to camouflage, but could also reflect an individual quality effect, maternal investment, or some other environmental effect. Our methods and results add to the toolbox for objective quantification of complex mammalian coat pattern traits, and should be useful for taxonomic or phenotypic classifications based on photographic coat pattern data.

Our analyses highlighted a few aspects of giraffe spots that were most likely to be heritable and which seem to have the greatest adaptive significance. Circularity and solidity, both descriptors of spot shape, showed the highest mother-offspring similarity. Circularity describes how close the spot is to a perfect circle, and is positively correlated with the trait of roundness and negatively correlated with aspect ratio. Solidity describes how smooth and entire the spot edges are versus tortuous, ruffled, lobed, or incised and is positively correlated with the trait of perimeter. We did not document significant similarity of any size-related spot traits (number of spots, area, perimeter, and maximum caliper), but the first dimension of the PCA was largely composed of size-related traits. These characteristics could form the basis for quantifying spot patterns of giraffes across Africa, and gives field workers studying any animal with complex color patterns a new quantitative lexicon for describing spots. However, our mode shade measurement was a crude metric, and color is greatly affected by lighting conditions, so we suggest standardization of photographic methods to control for lighting if color is to be analyzed in future studies.

We found that both size and shape of spots was relevant to fitness measured as juvenile survival. We observed the highest calf survival in the phenotypic group generally described as large spots that were either circular or irregular. Lowest survival was in the groups with small and medium-sized circular spots, and small irregular spots. Both the survival by phenotype analysis and the individual covariate survival analysis found that larger spots and irregularly shaped spots were correlated with increased survival. It seems likely that these traits enhance the background-matching of giraffe calves in the vegetation of our study area (Ruxton et al., 2004; Merilaita et al., 2017). However, covariation in spot patterns and survival could also reflect an individual quality effect, maternal investment, or some other environmental effect. The relationships among giraffe spot traits and their effects on fitness are clearly complex, and require additional investigations into adaptive function and genetic architecture.

Whether or not spot traits affect juvenile survival via anti-predation camouflage, spot traits may serve other adaptive functions such as thermoregulation (Skinner and Smithers 1990), social communication (VanderWaal et al., 2014), or indicators of individual quality (Ljetoff et al., 2007), and thus may demonstrate associations with other components of fitness, such as survivorship in older age classes or fecundity. Individual recognition, kin recognition, and inbreeding avoidance also could play a role in the evolution of spot patterns in giraffes (Beecher, 1982; Tibbetts & Dale, 2007; Sherman et al., 1997). Different aspects of spot traits may also be nonadaptive and serve no function, or spot patterns could be affected by pleiotropic selection on a gene that influences multiple traits (Lamoreux et al., 2010).

Photogrammetry to remotely measure animal traits has utilized geometric approaches that estimate trait sizes using laser range finders and known focal lengths (Lyon, 1994; Lee et al., 2016a), photographs of the traits together with a predetermined measurement unit (Ireland et al., 2006; Willisch et al., 2013), or lasers to project equidistant points on animals while they are photographed (Bergeron, 2007). We hope the framework we have described using ImageJ software to quantify spot characteristics with trait measurements from photographs will prove useful to future efforts at quantifying animal markings as in animal biometry (Kuhl & Burghardt, 2013). Trait measurements and cluster analysis such as we performed here could also be useful to classify subspecies, phenotypes, or other groups based on variation in markings, which could advance the field of phenomics for organisms with complex skin or coat patterns (Houle et al., 2010).

## Conclusions

Masai giraffe spot patterns are particularly diverse among giraffe subspecies (Dagg, 1968), and there are spot patterns in northern Tanzania that bear strong similarities to other giraffe subspecies elsewhere in Africa. Two recent genetic analyses of giraffe taxonomy both placed Masai giraffes as their own species (Brown et al., 2007; Fennessy et al., 2016), but the lack of quantitative tools to objectively analyze coat patterns for taxonomic classification may underlie some of the confusion that currently exists in giraffe systematics (Bercovitch et al., 2017). We expect the application of image analysis to giraffe coat patterns will provide a new, robust dataset to address taxonomic and evolutionary hypotheses.

Patterned coats of mammals are hypothesized to be formed by two distinct processes: a spatially oriented developmental mechanism that creates a species-specific pattern of skin cell differentiation and a pigmentation-oriented mechanism that uses information from the pre-established spatial pattern to regulate the synthesis of melanin (Eizirik et al., 2010). The giraffe skin has more extensive pigmentation and wider distribution of melanocytes than most other animals (Dimond & Montagna, 1976). Coat pattern variation may reflect discrete polymorphisms potentially related to life-history strategies, a continuous signal related to individual quality, or a combination of both. Future work on the genetics of coat patterns will hopefully shed light upon the mechanisms and consequences of coat pattern variation.

## Acknowledgements

This research was carried out with permission from the Tanzania Commission for Science and Technology (COSTECH), Tanzania National Parks (TANAPA), the Tanzania Wildlife Research Institute (TAWIRI), African Wildlife Foundation, and Manyara Ranch Conservancy. This paper was improved by comments from two anonymous reviewers and A.K. Lindholm.

